# Temporal discounting in adolescents and adults with Tourette syndrome

**DOI:** 10.1101/2020.02.14.947812

**Authors:** Canan Beate Schüller, Ben Jonathan Wagner, Thomas Schüller, Juan Carlos Baldermann, Daniel Huys, Julia Kerner auch Koerner, Eva Niessen, Alexander Münchau, Valerie Brandt, Jan Peters, Jens Kuhn

## Abstract

**Background:** Tourette syndrome is a neurodevelopmental disorder with the clinical hallmarks of motor and phonic tics which are associated with hyperactivity in dopaminergic networks. Dopaminergic hyperactivity in the basal ganglia has previously been linked to increased sensitivity to positive reinforcement and increases in choice impulsivity.

**Objective:** We address whether this extends to changes in temporal discounting, where impulsivity is operationalized as an increased preference to choose smaller-but-sooner over larger-but-later rewards. Results are discussed with respect to neural models of temporal discounting, dopaminergic alterations in Tourette syndrome and the developmental trajectory of temporal discounting.

**Methods:** In the first study we included nineteen adolescent patients with Tourette syndrome and nineteen age- and education matched controls. In the second study, we compared twenty-five adult patients with Tourette syndrome and twenty-five age- and education-matched controls.

**Results:** In the light of the dopaminergic hyperactivity model, we predicted differences in temporal discounting in patients with Tourette syndrome. However, computational modeling of choice behavior using hierarchical Bayesian parameter estimation revealed reduced impulsive choice in adolescent patients, and no group differences in adults.

**Conclusion:** We speculate that adolescents might show reduced discounting due to improved inhibitory functions that also affect choice impulsivity and/or the developmental trajectory of executive control functions. The absence of an effect in adults might be due to differences in the clinical population (e.g. patients who acquired successful tic inhibition during adolescence might have gone into remission). Future studies would benefit from adopting longitudinal approaches to further elucidate the developmental trajectory of these effects.

## 1. Introduction

Tourette syndrome (TS) is a childhood onset neuropsychiatric disorder characterized by motor and phonic tics that wax and wane in their severity with an estimated prevalence of around 1 % (1). Motor tics are repetitive, sudden movements such as eye blinking or facial muscle contractions and phonic tics are repetitive sounds such as throat clearing or verbal utterances (1,2). TS onset occurs predominantly in early childhood with a peak of symptom severity between the age of 10 and 12 years. Thereafter, tics improve in around 80 % of children until the end of adolescence (3,4). TS is associated with high comorbidity rates, predominantly attention-deficit/hyperactive disorder (ADHD), obsessive-compulsive disorder (OCD), depression (5) and impulse control disorders such as self-injurious behavior (6). Studies estimate that only 8 to 37 % of patients with TS do not exhibit any comorbidity (1,5,7). Treatment possibilities include cognitive behavioral therapy (i.e. habit reversal training) (8), antidopaminergic drugs (9) and new experimental approaches including cannabinoids (10) and deep brain stimulation (11,12).

Both clinical and neuroscientific research have highlighted possible developmental dysfunctions in the cortico-striatal-thalamo-cortical loops (13–15) especially with respect to dopamine (DA) that strongly modulates these circuits (16,17). The striatum, a main gateway in these loops (18) plays a key role in selectively amplifying converging sensory input to enable situation specific behavioral adaptations such as the adequate control of voluntarily movement (16). Predictions (i.e. expectations) of reward as well as the gating of specific motor responses are under dopaminergic modulation. Theories about the developmental underpinnings of TS in terms of DA function range from theoretical assumptions about a supersensitivity of striatal DA receptors (19) over tonic-phasic or presynaptic DA dysfunction (20–22) to DA hyperinnervation (20,23). Whereas the latter (i.e. excessive innervation of the basal ganglia via dendrites of midbrain DA neurons) may account for a range of empirical observations, including those, that led to the establishment of earlier hypotheses mentioned above (see 24).

To date several studies have investigated motor impulsivity in patients with TS with reference to DA’s role in reward and motor control (25,26). However, fewer studies have explored alterations in value-based decision-making in TS. However, this question is of particular interest because motor and choice impulsivity might at least in part be supported by common neural systems. First, DA in fronto-striatal circuits plays a role in both motor control (27,28) and choice impulsivity (29–33). Second, some studies have suggested that lateral prefrontal cortex regions might support impulse control functions, both in motor and non-motor domains (34–39). Two studies (40,41) examined impairments in value-based decision-making in TS in the context of reinforcement learning tasks. Palminteri and Pessiglione (2018) observed impaired learning from negative feedback in TS, which is consistent with the idea of a hyperdopaminergic state. Kéri and colleagues observed impaired probabilistic classification learning, especially in children with severe tics (41). However, whether choice impulsivity is impaired in TS remains an open question.

One way to reliably assess reward impulsivity (choice impulsivity) is via temporal discounting tasks (42–46). Temporal discounting describes a general preference for smaller sooner (SS) over larger, but later rewards (LL) (47). A relative preference for SS rewards (steep discounting of value over time) is associated with a range of problematic behaviors including substance use disorders and overweight/obesity (48) but also the tendency to procrastinate to invest for retirement (49) or to procrastinate to save up for future investments (50). The rate of temporal discounting is under complex modulation by individual and contextual variables (51–53), whereas striatal DA networks and prefrontal top down modulation seem to be the key regions of interest. However, the precise relationship between dopaminergic states and impulsive choice is complex. On the one hand, pharmacological reduction in DA levels decreases discounting (31–33,54). On the other hand, hyperdopaminergic states e.g. due to administration of the dopamine precursor L-DOPA, are also sometimes associated with increased discounting (29). Likewise, patients with Parkinson’s disease can exhibit increased impulsive behavior following DA replacement therapy (30). To sum up, DA modulation likely contributes to the modulation of intertemporal choice via its action on different fronto-striatal loops, but there is little evidence for a clear and simple linear relationship between DA levels and choice impulsivity.

In terms of top-down inhibitory mechanisms the picture is somehow relatively clear. The LPFC is assumed to modify choice impulsivity (55–58). That is, inhibition of the selection of tempting SS choices in this model depends on prefrontal inhibitory regulation of subcortical or ventromedial prefrontal value representations. Changes in structural and functional connectivity within this network are linked to the development of self-control (in this study the term self-control generally refers to far sighted behavior in value based decision making) from adolescence to early adulthood (59–61). Inhibition and top-down control likewise plays a central role in motor impulsivity and so is believed to modulate TS pathophysiology, e.g. in the context of suppressing urges and tics (25).

Studies did show that motor and cognitive impulsive actions might require different forms of the construct of self-control and can be differentiated (62). Even though it seems to play an important role in TS pathophysiology, evidence on the ability to successfully inhibit motor output in patients with TS is mixed and evidence is not entirely convincing that adolescents and adults show a general deficit in inhibitory control (25,26,63–67). However, there is extensive evidence for regional overlap between inhibitory mechanisms in terms of motor, choice impulsivity and even other forms like emotion regulation (35–38,68,69). Training in one domain might possibly affect performance other domains (70). Regarding choice and motor impulsivity the dorsal striatum might be a key region of interest where top down inhibitory processes (originating in PFC) modulate the execution or the re-evaluation of choice outcomes (71). These anatomical regions and attributed functions might be affected by TS pathophysiology (72)

To date it is still an open question whether patients with TS show aberrations in the domain of intertemporal choice. In the present study, we compared adolescents (Study 1, Hamburg) and adults (Study 2, Cologne) with TS to controls using two modified temporal discounting tasks. Based on the dopaminergic hyperinnervation model (24) we hypothesized that adolescents and adults with TS will show differences in temporal discounting compared to controls. We hope to broaden the understanding of value based decisions in TS on one operational measure of choice impulsivity that may predict, with unavoidable uncertainty, the vulnerability for short sighted behavior(49,50,73,74).

## 2. Methods and Materials

### 2.1 Ethics

The Ethics committee of the University of Cologne approved the study (protocol ID: DRKS00011748) and all participants provided written consent. Patients were recruited at the University Hospital of Cologne whereas healthy controls were recruited by advertisement. The Ethics committee of the University Hospital Hamburg approved the second study. Adolescents provided written assent and their parents provided written consent (PV4439). Adolescents with TS were recruited in the University Hospital of Hamburg and healthy adolescents were recruited by advertisement.

### 2.2 Study 1 methods (Adolescents)

#### 2.2.1 Participants

We included 19 adolescents with TS (mean(age): ± 14.21, SD: 2.37) and 19 age, education and gender-matched controls (mean(age): ± 14.21, SD: 2.53). All participants underwent a clinical assessment and performed a modified DD paradigm. Out of 19 adolescents, two were taking medication.

### 2.2.2 Clinical Assessment

Adolescents were assessed with the YGTSS (75), the PUTS (76) and the Children’s Yale-Brown Obsessive Compulsive Scale (CY-BOCS), a semi structured interview to evaluate OCD severity. For the CY-BOCS data are available from all the adolescents with TS and 13 controls; in total three adolescents with TS had a higher score than 12, which is an indicator for an OCD diagnosis (77). The “Fremdbeurteilungsbogen/Selbstbeurteilungsbogen für Aufmerksamkeitsdefizit-/Hyperaktivitätsstörungen” (FBB)-ADHD/(SBB)-ADHD is a diagnostic instrument to identify ADHD and includes a third-party assessment (FBB-ADHD) and self-reporting questionnaire (SBB-ADHS) (78). FBB-ADHD data is available for all adolescents with TS and 16 controls. SBB-ADHS data is available for 18 adolescents with TS and 17 controls. All adolescents also filled out a questionnaire on demographic measurements.

### 2.2.3 Temporal Discounting

The adolescents temporal discounting task consistent of 50 trials whereas patients and controls could choose between a SS reward (0, 1, 2, 3 or 4 cents) and a constant LL reward (5 cents), which was available after a varying waiting period (10, 20, 30, 40 or 60 seconds). The LL option was depicted with a blue circle and the SS option with a red circle both presented on a computer screen. The position of the red and blue circle was varied on a trial-wise manner whereas choice was indicated with a mouse click (see **Figure 1, supplemental data**). After each choice, the received reward (money) would be saved either immediately or after the appointed waiting period into a virtual saving account followed by visual feedback (displayed for 500ms). Thereafter, a blue screen with a black fixation cross was presented if the subject had chosen the immediate reward. The screen was presented for the same time the adolescents would have waited, had they chosen the delayed option (e.g. 20s if the waiting time for 5 cents would have been 20s). The overall task time was thereby kept constant, no matter whether participants chose predominantly SS or LL rewards. On a green bar below the choices, the participants could see how many trials had passed. Depending on the choices, participants could gain between 2.50 € and 5 € (79).

### 2.3 Study 2 methods (Adults)

#### 2.3.1 Participants

We recruited 25 patients (mean(age): ± 29.88, SD: 9.03) with TS diagnosed according to DSM-5 criteria (80) and 25 age, education and gender-matched controls (mean(age) ± 29.40, SD: 9.28). All participants underwent a clinical assessment, performed a temporal discounting paradigm, including a pretest based on prior procedures (see 81,82). Out of 25 patients, nine patients were taking medication or cannabinoids. Five patients were taking antidopaminergic drugs (Aripiprazole, risperidone, tiapride), one patient was taking an anticonvulsant (Orfiril) one patient was taking a noradrenergic and specific serotonergic antidepressant (Mirtazapine), and one patient was medicated with two antidopaminergic drugs (Aripiprazole, risperidone) and a selective serotonin reuptake inhibitor (Citalopram). One patient regularly smoked medical cannabis.

#### 2.3.2 Clinical Assessment

All participants performed on an equal clinical assessment. They filled out the Obsessive Compulsive Inventory-Revised (OCI-R) (83) and the Beck Depression Inventory (BDI) (84). The Wender Utah Rating Scale was used to assess ADHD symptoms (85). All participants filled out a short intelligence test (Leitprüfsystem-3 (LPS 3)) (86), followed by a demographic questionnaire with information on age, gender, handedness, years of education and current drug or alcohol use. Further, patients with TS completed an assessment with the Yale Global Tic Severity Scale (YGTTS) (75) and the premonitory urges scale (PUTS), a self-report scale to identify premonitory urges (76). All questionnaires were in German.

#### 2.3.3 Temporal discounting

##### Behavioral Pretest

All participants underwent an adaptive pretest to estimate an individual a-priori discount-rate via Maximum Likelihood estimation assuming and hyperbolic model (**Equation 2**, see below) and a softmax-Choice rule (**Equation 3**, see below). This rate was then used to create subject-specific trials for the following experimental session (see 81).

To create subject-specific trials, we used custom Matlab routines (MATLAB version 8.4.0. Natick, Massachusetts: The MathWorks Inc) (82). All experiments were administered using Presentation 16.3 (Presentation software; Neurobehavioral Systems, Inc).

##### Experimental session

The DD paradigm consisted of a series of 140 choices between direct (smaller-sooner (SS)) and delayed (larger-but-later (LL)) monetary rewards. The SS reward of 20€ was labeled as being available immediately whereas the LL reward was uniformly distributed (depending on the subject-specific pre-test) between 20.5€ and 80€ and available after 1, 2, 7, 14, 30, 90 or 180 days respectively. Participants were informed that one trial-choice combination would be selected at random and payed after the task was completed (see 85,86).

### 2.4 Analysis (Both studies)

#### Model free analysis

We first analyzed both datasets using model agnostic approaches to avoid possible caveats associated with model-based analysis, e.g., problems with parameter estimation or the choice for a theoretical framework (hyperbolic vs. exponential).

Due to task structure in study 1 (adolescents; see above), we used the percentage of LL in contrast to SS choices as a model agnostic quantification of choice behavior. For comparison we used a two-sided parametric test on the arc-sin-transformed values of SS vs. LL choices.

In study 2 (adults) we computed the area under the empirical discounting curve (*AUC*)(note, due to the number of varying delays, this procedure does not provide further information when applied to the data in study 1). In detail, the *AUC* corresponds to the area under the connected data points that describe the decrease of the subjective value (y-axis) over time (delay; x axis). Each specific delay was expressed as a proportion of the maximum delay and plotted against the normalized subjective (discounted) value. We then computed the area of the resulting trapezoids using **Equation 1**.

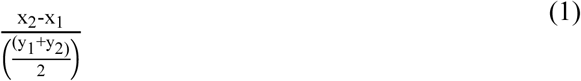

Smaller *AUC*-values indicate more discounting (more impulsive choices) and higher *AUC*-values indicate less discounting (less impulsive choices) (range between zero and one).

#### Computational modeling

Based on prior analysis and basic research in the field of temporal discounting we assumed a hyperbolic model (87,88) to describe the decrease in subjective value over time for both datasets (**Equation 2**). The LL reward that is delivered after a specific delay (D) is devaluated via a subject specific discount rate (*k*) that weights the influence of time on the subjective value (SV). A lower *k*-parameter reflects more patient preferences (reduced discounting) whereas a higher *k*-parameter reflects steeper discounting:

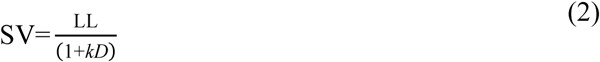

After devaluating the delayed option our model assumes that subjects compare the devaluated LL reward with the 20€ SS trial by trial and select the most valuable action under the influence of subject specific noise. This decision process between both subjective values is modeled by a simple softmax choice rule (**Equation 3**) where a free *temp* parameter scales the influence of value differences on choice. A high *temp* value implies that participants decide purely on value differences whereas lower values indicates higher choice stochasticity. For limit of temp=0 choices are random.

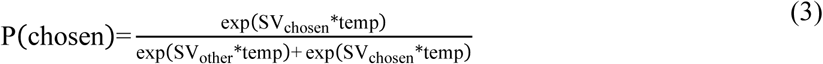

Models were fit using a hierarchical Bayesian framework to estimate parameter distributions via Markov Chain Monte Carlo (MCMC) sampling with Just Another Gibbs Sampler (JAGS) (89). Individual choice data were modeled using **Equations 2** and **3** (see above). Single subject parameters were drawn from group-level normal distributions, with mean and variance hyper-parameters that were themselves estimated from the data (see **Figure 1**). Model convergence was assessed via the RHAT statistic (Gelman-Rubinstein convergence diagnostic) where values < 1.01. (two chains) were considered acceptable.

**Figure 1.**
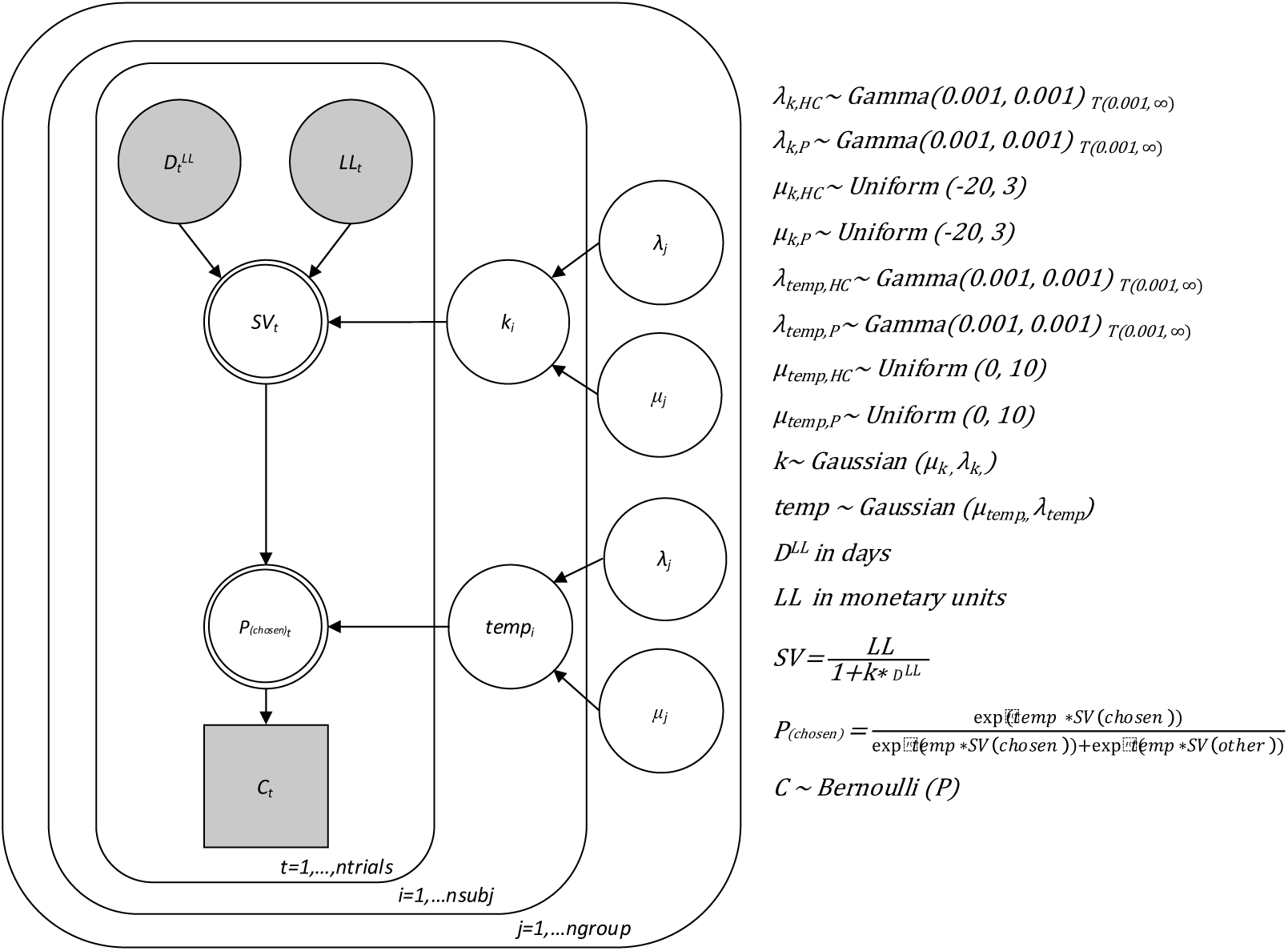
Hierarchical Bayesian model. Parameter estimates for each subject [*n* = 38 (study 1); *n* = 50 (study 2)], *k* (choice impulsivity) and *temp* (choice stochasticity) were drawn from different group distributions [ngroup = 2 (patients with TS (P)/ healthy controls (HC))] and mapped on the choice data [*n* = 50 (study 1); *n* = 140 (study 2)] for each participant.

**Figure 2.**
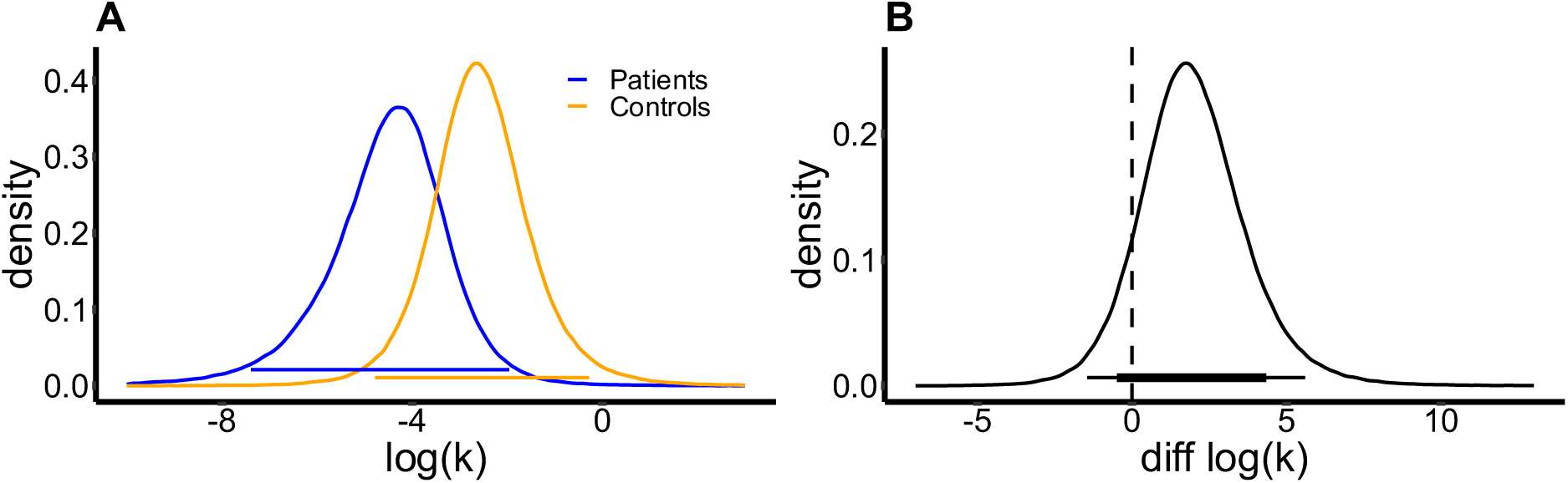
(A) Group level hyperparameter distributions of *log(k)* parameter for adolescents with TS (blue) and healthy controls (orange). (B) Difference distribution of (A) healthy controls minus adolescence with TS. The black bars indicate the 95% and 85% highest density intervals respectively. 89% of posterior hyperparameter samples exceed 0 (Note, even though 89% of samples exceed 0, the 85% HDI overlaps with 0. This is due to the fact that the HDI is computed differently from 10% and 90% quantiles or the subtraction of posterior samples which are just in Bayesian comparisons of posterior distributions (see Kruschke 2011 for details).In other words 89% of the target distribution from adolescents with TS is lower than the equivalent distribution in healthy controls. This can be interpreted as a chance of 89% of decreased discounting of delayed rewards in adolescents with TS when compared to healthy controls.

**Figure 3.**
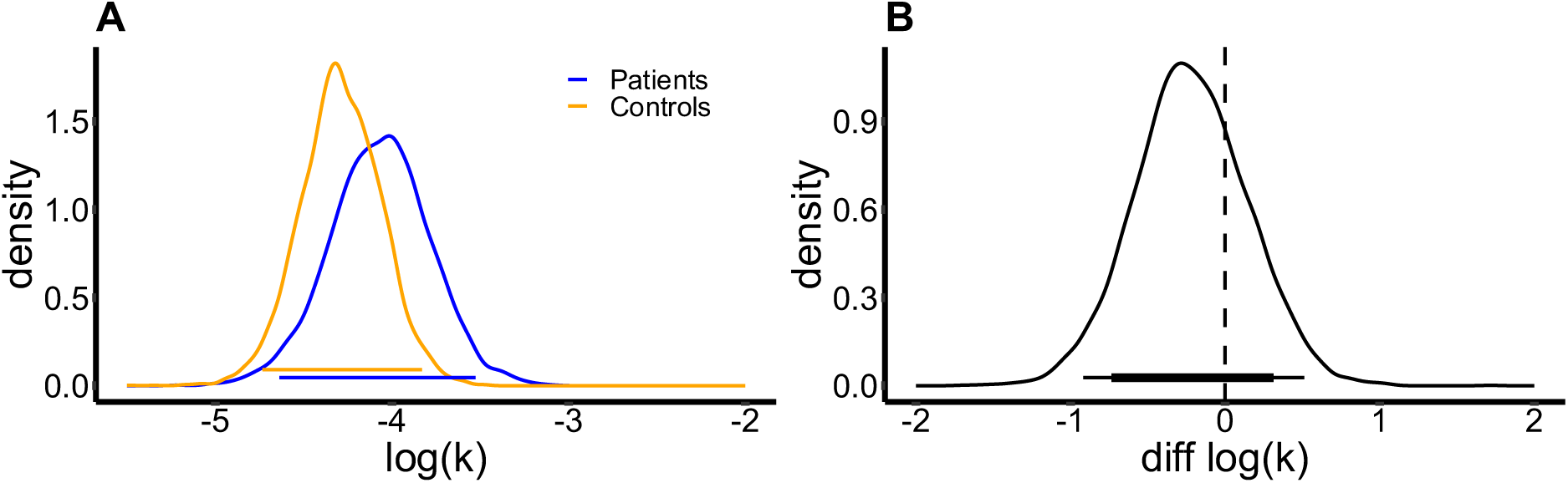
(A) Group level hyperparameter distributions of *log(k)* for patients with TS (blue) and healthy controls (orange); The orange and blue bars indicate the 95% highest density interval for each group (B) Difference distribution of hyperparameters shown in (A) - healthy controls minus patients with TS. The black bars indicate the 95% and 85 % highest density interval respectively. 72 % of the hyperparameter distribution is below 0 which can be interpreted as a chance of 72% of steeper discounting in patients with TS.

Group comparisons were conducted by examining the differences in posterior distributions per parameter of interest. The strength of evidence for directional effects was examined by computing directional Bayes Factors for each group level difference distribution. A Bayes factor > 3 yields positive evidence (90).

## 3. Results

### 3.1 Study 1 (Adolescents)

#### 3.1.1 Demographic characteristics and clinical assessment

Demographic and clinical characteristics between adolescents with TS and controls are shown in **Table** For demographic, clinical and neuropsychological characteristics of adolescents with TS and controls adjusted for multiple comparison see **Table 1, supplemental data**.

#### 3.1.2 Temporal discounting

##### Model free analysis

Controls chose the LL reward in 48.3 % of all cases whereas adolescents with TS chose the that option in 58.4 % of all trials (see **Figure 2, supplemental data**). Before using a parametric-test we applied an arcsin-transformation on all mean choice proportions (by participant) and then tested for group differences. Even though patients with TS did choose the LL option around 10% more often both groups did not differ significantly (*T* =1.0646; df = 35.83; *p* = 0.29).

##### Computational modeling

Examination of the posterior distributions of log(*k*) from the computational model revealed attenuated impulsive choice (smaller log(*k*)) in patients with TS: 89.07 % of the log (*k*) posterior difference distribution (controls *minus* patients) exceeded 0, suggesting steeper discounting of value over time in controls (**Equation 2**). Computing a directional Bayes Factor (dBF) for the group difference yielded dBF=8.13, that is, given the data, a reduction in discounting in patients was 8.13 times more likely than an increase. For analysis of choice stochasticity see **Figure 4, supplemental data**.

### 3.2 Study 2 (Adults)

#### 3.2.1 Demographic characteristics and clinical assessment

Demographic and clinical characteristics of patients with TS and controls are shown in **Table 2**. Controls did not score in any clinically relevant ranges. Neither patients nor controls reported clinically relevant drug or alcohol abuse.

**Table 1.** Demographic, clinical and neuropsychological characteristics of adolescents with TS and healthy controls.

|  | Adolescents with TS<br>(n=19) |  | Healthy controls<br>(n=19) |  | *T/U/ $\chi^2$* | *p* |
| --- | --- | --- | --- | --- | --- | --- |
|  | Mean | SD | Mean | SD |  |  |
| Age (Years) <sup>a</sup> | 14.21 | 2.37 | 14.21 | 2.53 | 0.000 | 1.000 |
| Male/Female <sup>c</sup> | 13/6 | - | 78.9 | - | 0.543 | 0.467 |
| Right-handed <sup>c</sup> | 14/19 | - | 84.2 | - | 1.276 | 0.435 |
| Current medication | 2/19 | - | - | - | - | - |
| YGTSS impairment | 16.00 | 8.00 | - | - | - | - |
| YGTSS | 23.37 | 12.38 | - | - | - | - |
| PUTS | 19.53 | 5.61 | - | - | - | - |
| FBB-ADHD<sup>b</sup> | 0.38 | 0.26 | 0.82 | 0.48 | -3.226 | 0.093 |
| SBB-ADHD<sup>a</sup> | 0.39 | 0.22 | 0.68 | 0.39 | 88.0 | 0.497 |
| CY-BOCS <sup>b</sup> | 6.84 | 6.31 | 0.08 | 0.277 | 21.50 | <0.001 |

ADHD, attention deficit hyperactivity disorder; CY-BOCS, Children's Yale-Brown Obsessive-Compulsive Scale; (FBB)-ADHD/(SBB)-ADHD, Fremdbeurteilungsbogen/Selbstbeurteilungsbogen für Aufmerksamkeitsdefizit-/Hyperaktivitätsstörungen; PUTS, Premonitory Urge for Tics Scale; TS, Tourette syndrome; YGTSS, Yale Global Tic Severity Scale.
a. T-test was used because data was normally distributed.
b. Mann Whitney U test was used because data was not normally distributed.
c. $\chi^2$ square test.

**Table 2:**
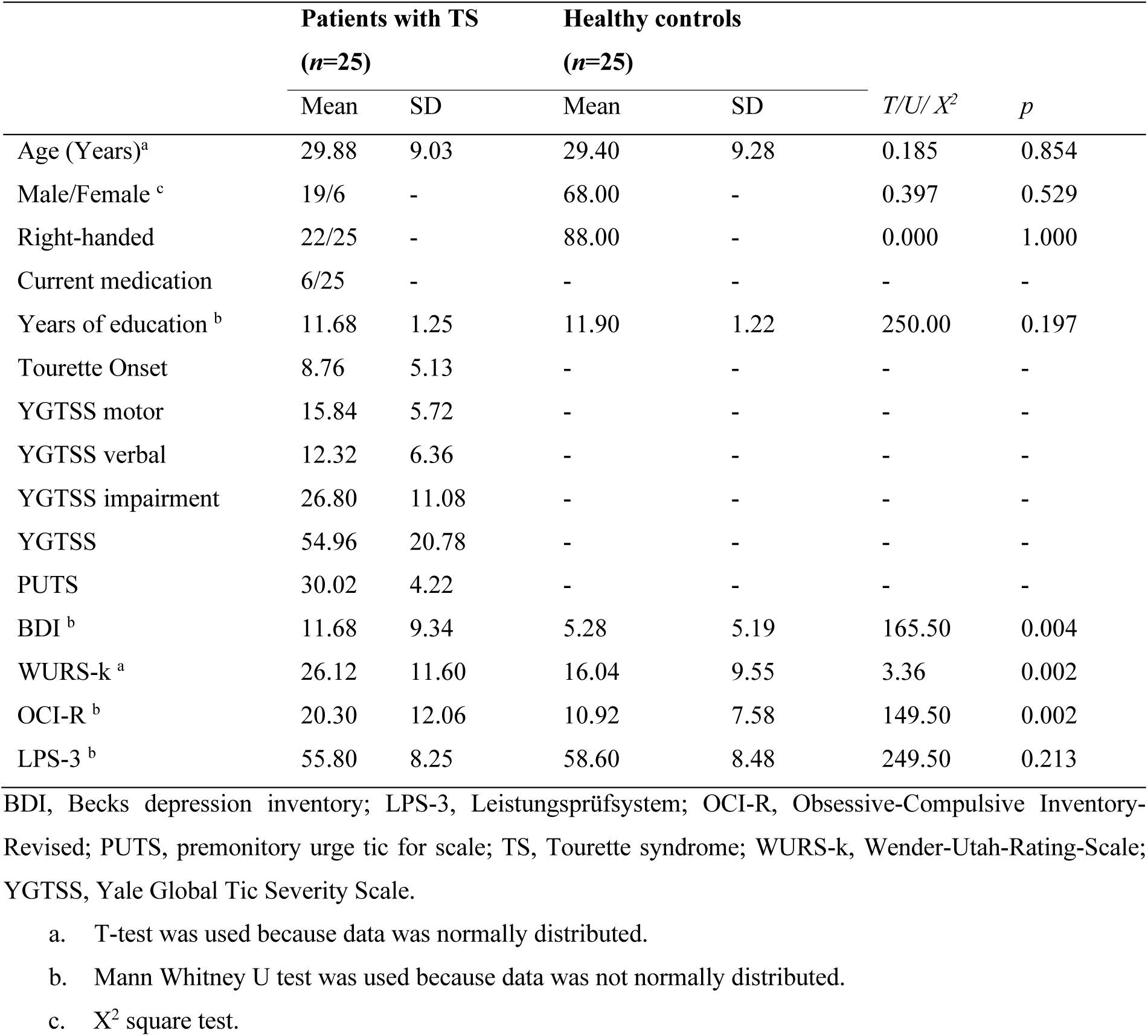
Demographic, clinical and neuropsychological characteristics of patients with TS and healthy controls.

|  | Patients with TS<br>(*n*=25) |  | Healthy controls<br>(*n*=25) |  | *T/U/ X*<sup>2</sup> | *p* |
| --- | --- | --- | --- | --- | --- | --- |
|  | Mean | SD | Mean | SD |  |  |
| Age (Years)<sup>a</sup> | 29.88 | 9.03 | 29.40 | 9.28 | 0.185 | 0.854 |
| Male/Female <sup>c</sup> | 19/6 | - | 68.00 | - | 0.397 | 0.529 |
| Right-handed | 22/25 | - | 88.00 | - | 0.000 | 1.000 |
| Current medication | 6/25 | - | - | - | - | - |
| Years of education <sup>b</sup> | 11.68 | 1.25 | 11.90 | 1.22 | 250.00 | 0.197 |
| Tourette Onset | 8.76 | 5.13 | - | - | - | - |
| YGTSS motor | 15.84 | 5.72 | - | - | - | - |
| YGTSS verbal | 12.32 | 6.36 | - | - | - | - |
| YGTSS impairment | 26.80 | 11.08 | - | - | - | - |
| YGTSS | 54.96 | 20.78 | - | - | - | - |
| PUTS | 30.02 | 4.22 | - | - | - | - |
| BDI <sup>b</sup> | 11.68 | 9.34 | 5.28 | 5.19 | 165.50 | 0.004 |
| WURS-k <sup>a</sup> | 26.12 | 11.60 | 16.04 | 9.55 | 3.36 | 0.002 |
| OCI-R <sup>b</sup> | 20.30 | 12.06 | 10.92 | 7.58 | 149.50 | 0.002 |
| LPS-3 <sup>b</sup> | 55.80 | 8.25 | 58.60 | 8.48 | 249.50 | 0.213 |

BDI, Becks depression inventory; LPS-3, Leistungsprüfsystem; OCI-R, Obsessive-Compulsive Inventory-Revised; PUTS, premonitory urge tic for scale; TS, Tourette syndrome; WURS-k, Wender-Utah-Rating-Scale; YGTSS, Yale Global Tic Severity Scale.
a. T-test was used because data was normally distributed.
b. Mann Whitney U test was used because data was not normally distributed.
c. X<sup>2</sup> square test.

#### 3.2.2 Temporal Discounting

##### Model free analysis

Applying a parametric t-test on the integral of the area under the empirical discounting curve revealed no significant differences between patients with TS (mean(*AUC*) = 0.459) and controls (mean(*AUC*) = 0.511) (*t* = −0.8791; *df* = 46.1; *p* = 0.38), see **Figure 3, supplemental data**.

##### Computational modeling

In line with our model-agnostic approach examination of the posterior distributions of log(*k*) from the computational model revealed only minor differences in impulsive choice (smaller log(*k*)) in controls as only 38 % of the log (*k*) posterior difference distribution (controls *minus* patients) exceeded 0. This suggests no significant differences in discounting of value over time. Computing a directional Bayes Factor (dBF) for the group difference yielded dBF=0.38 (no mentionable evidence), that is, if anything a descriptive decrease in discounting in controls when compared to patients with TS. In consequence absolute log(*k*) distributions showed substantial overlap between groups (**Figure 4A**). Since some patients with TS were treated with antidopaminergic drugs we excluded these six subjects and repeated our computational analysis. The exclusion of these patients only had marginal effects and the result pattern did not change. For analysis of choice stochasticity see **Figure 5, supplemental data**.

Further, no ties between subject specific choice impulsivity (median(*k*)) and choice stochasticity (median(*temp*)) parameters and our questionnaire data could be detected. In detail, we performed a simple correlation analysis (corrected for multiple comparisons; note an additional exploratory analysis without correcting for multiple corrections did not reveal any significant correlation) in between the before mentioned model parameters and the following inventories: WURSK-k scale, OCI-R and BDI (see **Table 2, supplemental data**).

## 4. Discussion

We examined temporal discounting in adolescents and adults with TS. Based on neural models of the etiology of TS we predicted increased temporal discounting in TS due to a putative increase in DA signaling (23). In contrast to our prediction, computational modeling using hierarchical Bayesian parameter estimation revealed that adolescent TS patients showed reduced temporal discounting compared to controls. In contrast, we observed little evidence for robust group differences in adult TS patients.

TS is a complex neuropsychiatric disorder that is associated with developmental dopaminergic anomalies and a failure to control involuntary actions (1,2,24–26). These dopaminergic anomalies may either cause, enable or enhance tics via inadequate gating of information through the striatum (16). In the current study, we report data from two temporal discounting tasks to examine if self-controlled choices are under modulation of TS pathophysiology. We hypothesized that dopaminergic anomalies might interfere with the valuation of decision options, which are modulated by both dopaminergic signaling and prefrontal inhibitory control (24–26).

The DA hyperinnervation model of TS, in conjunction with some of the empirical findings linking elevated DA to increased human temporal discounting (24,29), might then predict increased discounting in patients with TS. However, other studies point towards reductions in temporal discounting due to pharmacological elevation of DA levels. Generally, the human literature on dopaminergic contributions to impulsivity is characterized by substantial heterogeneity (91). A further complicating factor is that dopaminergic effects might be non-linear (92), as summarized in the inverted U-model of DA functioning (93). However, it is obvious that transient pharmacological dopaminergic interventions in healthy subjects and long-term abnormal dopaminergic states in neurodevelopmental conditions such as TS will have markedly different behavioral effects. Nevertheless, our results suggest that the putative chronic hyperdopaminergic state of TS does not give rise to substantial changes in temporal discounting in adults.

In contrast, we did find evidence for a moderate decrease in temporal discounting in adolescents with TS when compared to healthy controls. Our analysis revealed that a decrease in temporal discounting in adolescents with TS was about 8 times more likely than an increase (dBF = 8.13). Adolescents typically show higher discount rates than adults (94,95). This is thought to be attributable to increases in functional and structural fronto-subcortical connectivity that continue until early adulthood (26,59–61). Adolescents with TS are constantly faced by tics and the need to control their motor output. Even though these tics might emerge from complex neurophysiological interactions i.e. hyperactive DA modulated striatal gating and reduced inhibition of GABAergic interneurons (96,97), one could speculate that the ability to inhibit tics might foster the ability to inhibit other impulses thereby strengthening cognitive control more generally (70). The question then arises why such an effect would not likewise translate into greater self-control during temporal discounting in the adult TS patients as well. One possibility is that such a “training” account merely affects the developmental trajectory of self-control, such that adolescents with TS reach adult levels of self-control earlier than their healthy peers. Testing such a model would of course require longitudinal studies.

Additional clinical differences between adolescent and adult TS patients further complicate the interpretation of the differential effects in the two age groups. Adolescents and adults with TS exhibit different tic-phenomenology, adolescents exhibit less variability and/or fluctuations in tics as well as additional comorbidities such as autistic spectrum disorders and oppositional defiant disorder (1). Adolescents who successfully control their tics have a greater likelihood of eventual remission, likely due to better executive control capabilities (98). In contrast, patients who still exhibit TS in adult life exhibit attenuated inhibitory control (66). In both samples, the discount rate (*k*) was not significantly correlated (corrected for multiple comparisons) with ADHD, OCD comorbid symptomatology or the YGTSS (see **Table 1** and **Table 2, supplemental data**).

The present study has several limitations. First, adolescents and adults performed different temporal discounting tasks with different reward magnitudes (0-4 cents vs. 20-80€) on a different timescale (immediate up to a minute (adolescents) vs. immediately after the task to up to weeks (adults)). Reward magnitudes in the range of cents vs. tens of Euros may entail different valuation and/or control processes (99,100). This precludes direct comparisons in log(*k*) between age groups. Second, we do draw theoretical conclusions from reward impulsivity to motor inhibition in patients with TS, even though we do not compare motor inhibition empirically. Further studies should try to further examine the developmental trajectories of both of them. Third, although only two adolescents with TS took medication, about a quarter of adult patients (*n*=6) were on antipsychotic medication. An integrative review showed that most TS medication (i.e. D_2_ antagonists) reduce phasic DA, tonic DA or both (24) and DA dysfunction in cortico-striatal-thalamo-cortical was likely affected by the medication. However, a control analysis in which all medicated participants were excluded yielded the same pattern of results. Finally, patients and controls in the two different studies did not complete the exact same set of questionnaires (i.e. LPS-3).

## 5. Conclusion

The present study assessed temporal discounting in adolescent and adult TS patients, as well as matched healthy controls. Our data suggest reduced discounting in adolescent TS patients compared to matched controls. We speculate that this might be due to improved inhibitory functions that affect choice impulsivity and/or the developmental trajectory of executive control functions. Interestingly, adult patients with TS exhibited levels of discounting similar to controls. This might be due higher disease severity in adult patients with TS (e.g., patients who acquired successful tic inhibition during adolescence might have gone into remission). Future studies would benefit from adopting a longitudinal approach to further elucidate the developmental trajectory of these effects, and from directly examining effects of dopaminergic medication on these processes in TS.

## Acknowledgments

We would like to thank Milena Marx for support in Figure preparation.

## Authors Roles

Canan Beate Schüller: 1A, 1B, 1C, 2B, 3A; Ben Jonathan Wagner: 2A, 2B, 3A; Thomas Schüller: 1A, 3B; Juan Carlos Baldermann 1A, 3B:, Daniel Huys:1A, 3B; Julia Kerner auch Koerner:1C, 1B, 3B; Eva Niessen:1A, 1C, 3B; Alexander Münchau: 1A, 3B, Valerie Brandt: 1A, 1B, 1C, 3B; Jan Peters, 1A, 1C, 2A, 2C, 3B:, Jens Kuhn: 1A, 3B.

## Financial Disclosures for all authors (for the preceding 12 months)

Canan Beate Schüller: Walter and Marga Boll Foundation; Ben Jonathan Wagner: None; Thomas Schüller: Walter and Marga Boll Foundation; Juan Carlos Baldermann: None; Daniel Huys: None; Julia Kerner auch Koerner: None; Eva Niessen: None; Alexander Münchau: Commercial research support: Pharm Allergan, Ipsen, Merz Pharmaceuticals, Actelion; Honoraria for lectures: Pharm Allergan, Ipsen, Merz Pharmaceuticals, Actelion, GlaxoSmithKline, Desitin and Teva; Support from Foundations: Possehl-Stiftung (Lübeck, Germany), Margot und Jürgen Wessel Stiftung (Lübeck, Germany), Tourette Syndrome Association (Germany), Interessenverband Tourette Syndrom (Germany), CHDI; Academic research support: Deutsche Forschungsgemeinschaft (DFG): projects 1692/3-1, 4-1, SFB 936, and FOR 2698 (project numbers 396914663, 396577296, 396474989), Innovationsausschuss of the Gemeinsamer Bundesausschuss: Translate NAMSE (structural support for the Lübeck Center for Rare Diseases); Royalties for the book Neurogenetics (Oxford University Press); Advisory Boards: German Tourette syndrome association; Alliance of patients with chronic rare diseases; Valerie Brandt: Public Engagement with Research Fund, the ROLI music company, Tourettes Action; Jan Peters: Grants: DFG PE1627/5-1 und PE1627/5-2; Jens Kuhn: received honoraria from Bayer, Janssen, Lundbeck, Neuraxpharm, Otsuka Pharma, Schwabe and Servier for lecturing at conferences and financial support to travel. He has received financial support for Investigator initiated trials from Medtronic GmbH.

## Supplemental data

### Tables

**Table 1.** Demographic, clinical and neuropsychological characteristics of adolescents with TS and healthy controls adjusted for multiple comparison (using holm’s method; note: an additional exploratory analysis without correcting for multiple corrections did not reveal any significant correlation).

| Questionnaire/<br>subscale | Patients with TS (*n*=19) |  | Healthy controls (*n*=18) |  |
| --- | --- | --- | --- | --- |
|  | median(*k*)/<br>*r* (*p*-value) | median(*temp*)/<br>*r* (*p*-value) | median(*k*)/<br>*r* (*p*-value) | median(*temp*)/<br>*r* (*p*-value) |
| SBB Attentional | 0.25 (0.32[1.00]) | 0.03 (0.90[1.00]) | -0.02 (0.94[1.00]) | -0.02 (0.93[1.00]) |
| SBB_Motor | 0.33 (0.18[1.00]) | -0.04 (0.87[1.00]) | 0.20 (0.44[1.00]) | 0.00 (0.99[1.00]) |
| SBB Impulsive | -0.17 (0.50[1.00]) | 0.43 (0.07[0.71]) | 0.24 (0.35[1.00]) | -0.12 (0.63[1.00]) |
| SBB ADHD | 0.12 (0.65[1.00]) | 0.22 (0.37[1.00]) | 0.13 (0.60[1.00]) | -0.05 (0.85[1.00]) |

**Table 2.** Correlation analysis in adult patients with TS and healthy controls adjusted for multiple comparison (using holm’s method).

| Questionnaire/<br>subscale | Patients with TS (*n*=25) |  | Healthy controls (*n*=25) |  |
| --- | --- | --- | --- | --- |
|  | median(*k*)/<br>*r* (*p*-value) | median(*temp*)/<br>*r* (*p*-value) | median(*k*)/<br>*r* (*p*-value) | median(*temp*)/<br>*r* (*p*-value) |
| WURSK-k | -0.03 (1.00) | -0.14 (1.00) | 0.29 (1.00) | 0.09 (1.00) |
| OCI-R | 0.20 (1.00) | -0.39 (1.00) | 0.31 (1.00) | -0.20 (1.00) |
| BDI | -0.18 (1.00) | 0.43 (1.00) * | 0.28 (1.00) | 0.03 (1.00) |

BDI, Becks depression inventory; OCI-R, Obsessive-Compulsive Inventory-Revised; TS, Tourette syndrome; WURSK-k, Wender-Utah-Rating-Scale.

### Figures

**Figure 1:**
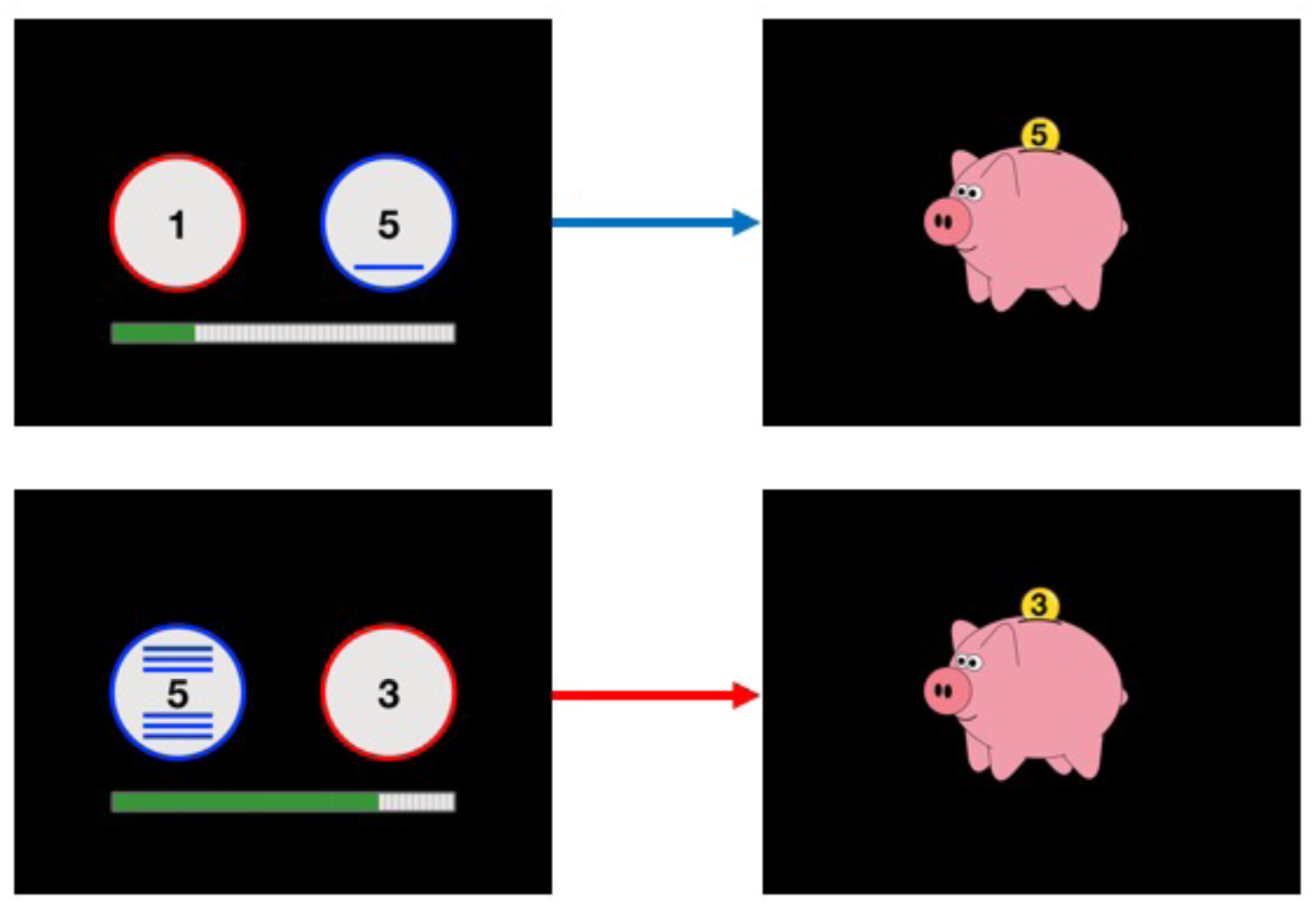
Example for two trials in the temporal discounting task adapted for children and adolescents. The blue circle shows the reward (in cents) that the participant will receive if they wait. How long they have to wait is indicated by the lines, i.e. one blue line = 10s wait, 6 blue lines = 60s wait. The red circle indicates how much the participant will receive if they move on to the next trial immediately (0-4 cents). Participants received feedback about the amount earned after every trial (piggy bank). The green bar below the two circles indicates how many trials the participant has already finished.

**Figure 2.**
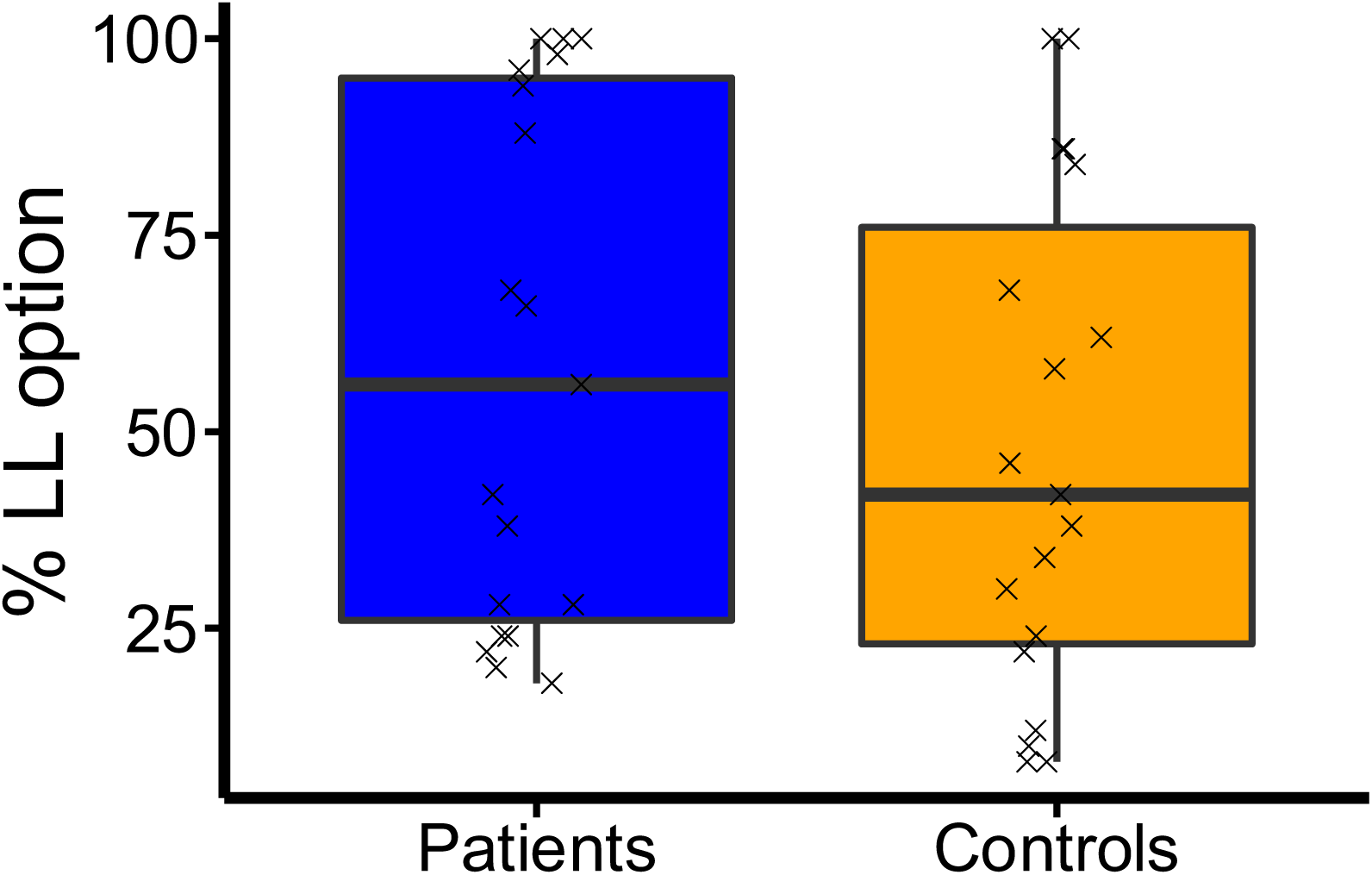
Percentage of larger, but later (LL) choices in adolescents with TS and healthy controls.

**Figure 3.**
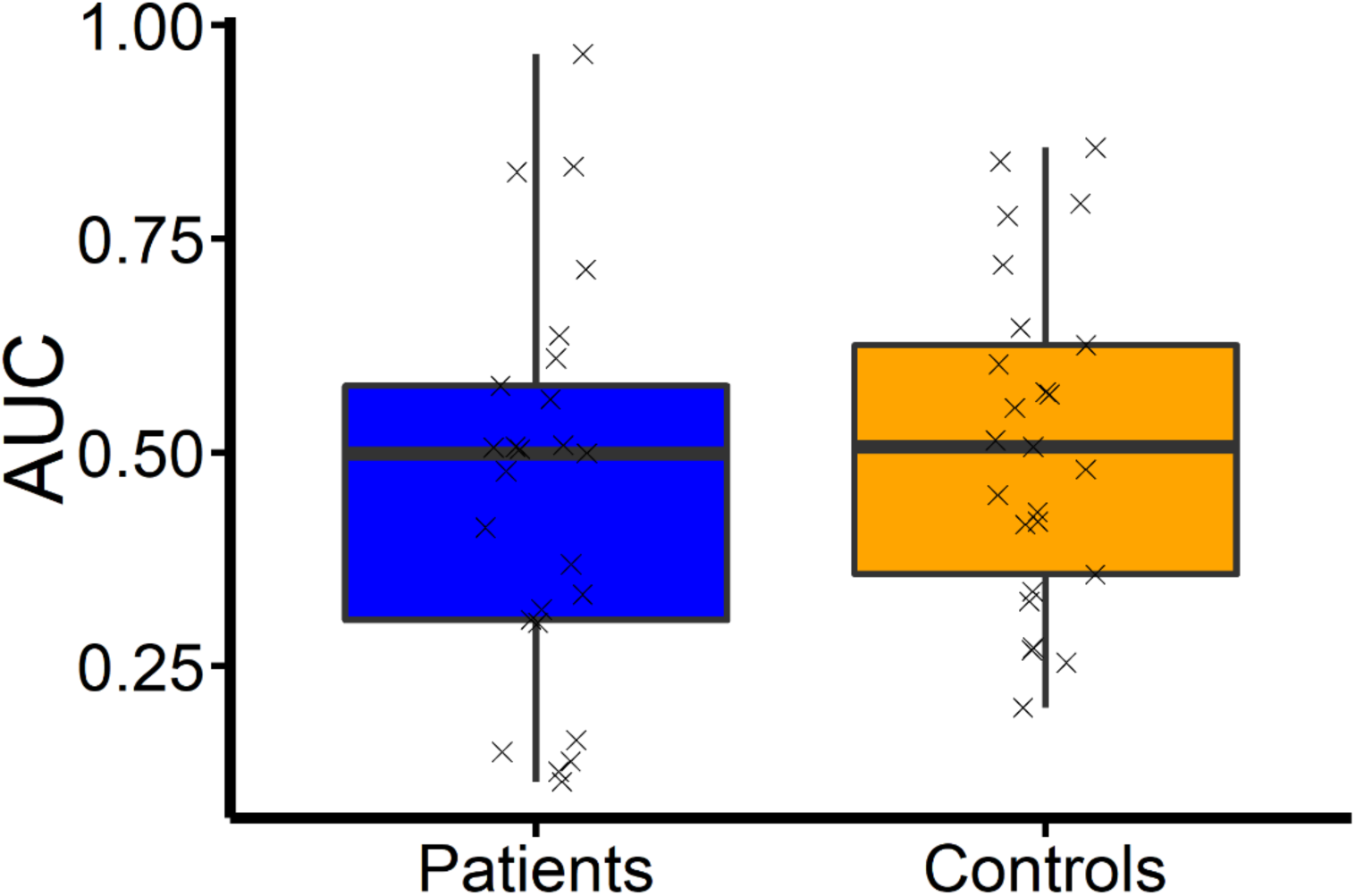
Subject specific measurements of the integral under the empirical area under the curve in adults with TS and healthy controls.

### Study 2 (adolescent)

#### Choice stochasticity

We applied the identical as for log(*k*) to the inverse temperature parameter (see **Equation 3**) and yielded a dBF of 0.66 (no mentionable evidence) implying no difference in between controls and adolescent patients with TS concerning value independent noisy choices.

**Figure 4.**
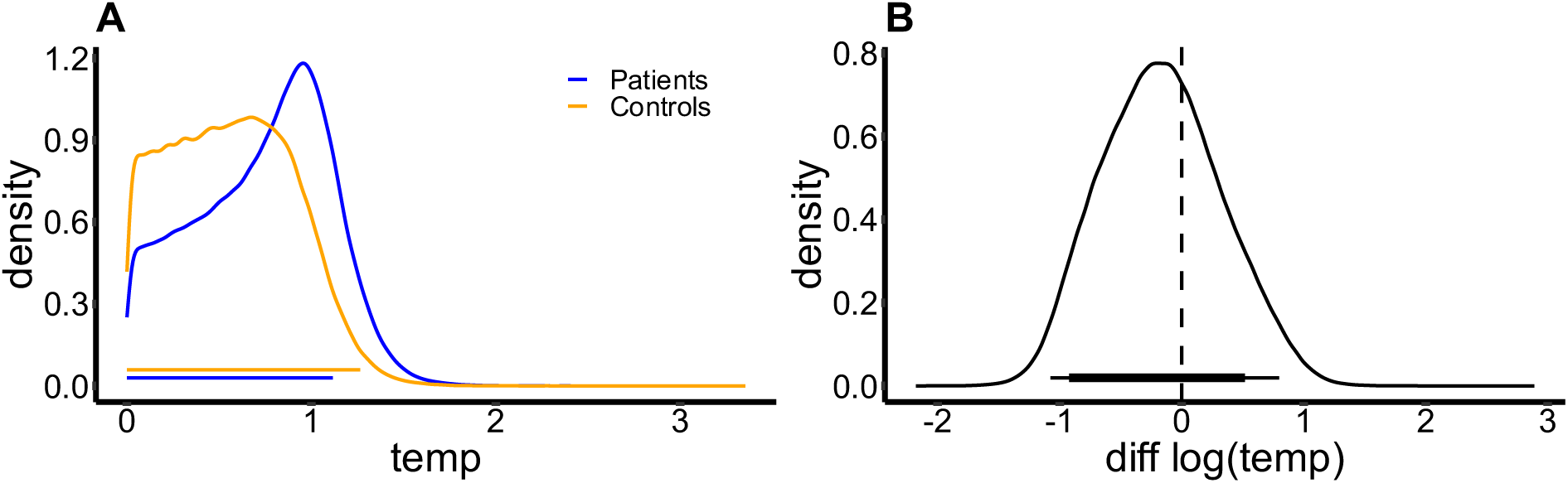
**(A)** Group-level hyperparameter distributions of the decision noise parameter temp for adolescents with TS (blue) and healthy controls (orange). **(B)** Difference distribution of temp hyperparameter healthy controls minus adolescents with TS.

### Study 2 (adults)

#### Choice stochasticity

The additional analysis for choice stochasticity yielded a dBF of 0.67 indicating no substantial difference in decision noise. We observed a substantially higher variance in the decision noise (*temp*) hyperparameter in controls, an effect that was driven by a few participants with very high and some with very low decision noise. In contrast, in adult patients with TS, observed temp values were more homogenous.

**Figure 5.**
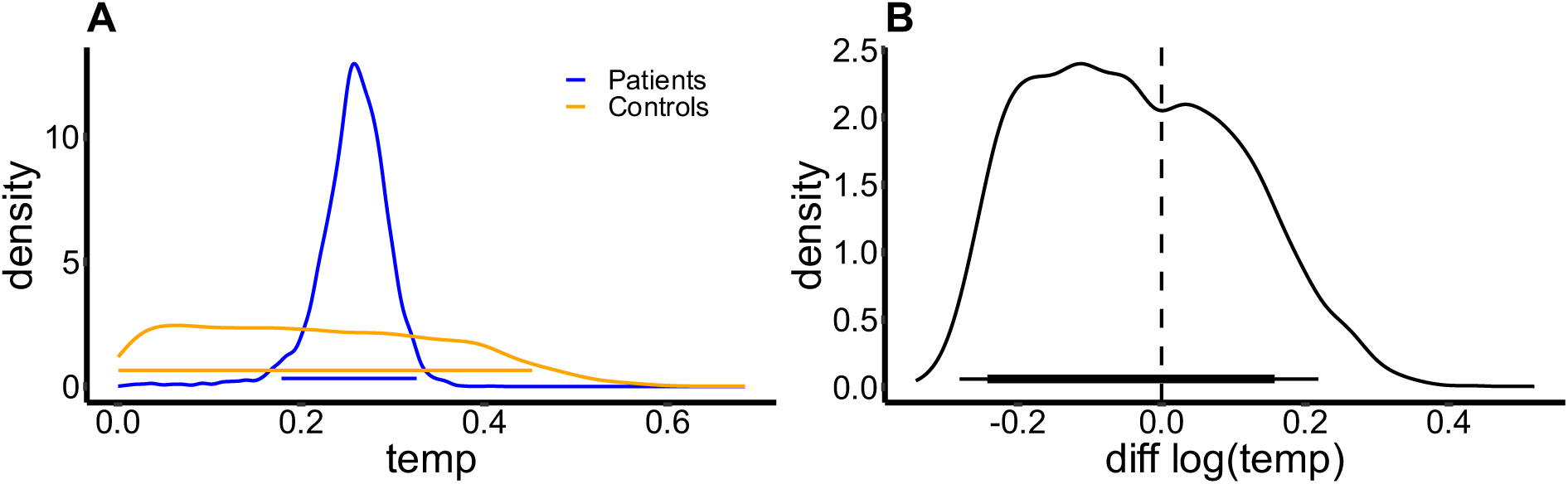
(**A**) Group-level hyperparameter distributions of the decision noise parameter temp for TS patients and healthy controls. **(B)** Difference distribution of temp hyperparameter TS patients – healthy controls with 95% and 85% highest density intervals.

## Notes

**Funding sources** C.S. and T.S. were funded by the Walter and Marga Boll Foundation. J.P. was supported by Deutsche Forschungsgemeinschaft (DFG: PE 1627/5-1 and PE 1627/5-2). A.M. was supported by the DFG (FOR 2698).

